# CDEK: Clinical Drug Experience Knowledgebase

**DOI:** 10.1101/474189

**Authors:** Rebekah H. Griesenauer, Constantino Schillebeeckx, Michael S. Kinch

## Abstract

The Clinical Drug Experience Knowledgebase (CDEK) is a database and web platform of active pharmaceutical ingredients with evidence of clinical testing as well as the organizations involved in their research and development. CDEK was curated by disambiguating intervention and organization names from ClinicalTrials.gov and cross-referencing these entries with other prominent drug databases. Approximately 43% of active pharmaceutical ingredients in the CDEK database were sourced from ClinicalTrials.gov and cannot be found in any other prominent compound-oriented database. The contents of CDEK are structured around three pillars: active pharmaceutical ingredients (n = 22,292), clinical trials (n = 127,223), and organizations (n = 24,728). The envisioned use of the CDEK is to support the investigation of many aspects of drug development, including discovery, repurposing opportunities, chemo- and bio-informatics, clinical and translational research, and regulatory sciences.

Database URL: http://cdek.wustl.edu

## INTRODUCTION

The process in which drugs are discovered and developed has fundamentally changed since the inception of the pharmaceutical industry and continues to evolve. Several research groups have peered into the past to identify trends in pharmaceutical innovation based upon FDA approved medicines (1–3). The Center for Research Innovation in Biotechnology (CRIB) at Washington University in St. Louis is amidst an ongoing effort to objectively track and analyze trends in the innovation of new medicines. Several published works were facilitated by analysis of a precursor database (curated and maintained by CRIB) of all FDA-approved new molecular entities (NMEs), which included their mechanistic basis, therapeutic applications, and organizations guiding their clinical development. This NME database (http://cribdb.wustl.edu) also includes products that were once approved but no longer marketed as a result of toxicity, lack of efficacy, obsolescence, production issues, or lack of demand.

A handful of reviews on the biopharmaceutical industry trends and innovation sources revealed a trove of findings, many unexpected, and all supported by objective data (all of which we have made public). As one example, a handful of organizations have recently come to control two-thirds of NMEs and these marketing organizations often have little or no internal drug discovery or development activities (4). Whereas large, traditional pharmaceutical companies receive most FDA approvals, upstart biotechnology companies increasingly dominate early-stage discovery (including patents and Investigational New Drug (IND) applications) (5). The NME database also revealed the causes and impact of corporate consolidation in transforming research and development. Whereas 60% of all acquired biotechnology companies were acquired within 5 years (before or after) their first NME approval was granted, the number of new organizations to receive their first approval has not kept pace (6). Consequently, the net number of research organizations that remain active and independent in new drug research has has eroded from over 200 firms in 2004 to 100 firms at the end of 2015 (7).

Based on findings with FDA-approved medicines, we analyzed the mechanistic basis and therapeutic indications of FDA approved medicines and changes over time. In some cases, these works emphasized therapeutic areas (e.g., the decline in anti-infectives or the rise in oncology (8)) while others focused upon drug targets, revealing three target families dominate FDA-approved drugs (G-protein coupled receptors, membrane channels and transporters, and targets involving nuclear signaling (9)). Beyond clinical indications and drug targets, exploration of other facets of the biotechnology industry enabled by the NME database included regulatory pathways and timelines (10), vaccine development (11), and the rise of biologics (12).

Although intriguing, we considered prior observations of pharmaceutical research and development trends to be undoubtedly skewed by focusing only upon FDA-approved medicines. It is generally understood most drug research does not conclude with a single FDA-approval as post-approval research (e.g., additional indications or post-approval commitments) capture an ever-increasing fraction of research and development expenditures and are not captured in analyses of drugs based solely upon a designation of “FDA-approved.” Compounding the problem, the timelines required for drug development mean an FDA-approval reflects research and development activities that were likely initiated more than a decade before, enfeebling any analyses intended to assess current or predict future research and development activity. Consequently, conjectures and definitive conclusions are not feasible absent a more comprehensive accounting of drug development efforts; including an assessment of successes, failures, and those experimental medicines currently being developed.

Powerful insights can be obtained by analyzing and modeling drug “failures.” In Gayvert et al. (13), a random forest machine learning algorithm classified a set of compounds as “FDA approved” or “Failed for Toxicity” based on chemical structure and drug target features. In this study, 784 FDA approved drugs and 100 “toxic” drugs were used to train and validate the machine learning model. Ideally, failed drugs would have made up a higher percentage of the sample, but sufficient data on failed drugs are not readily available. Nonetheless, these findings revealed machine learning predictions can be quite powerful provided that they are supplied with enough data for training and validation. Wong et al. (14) were able to assign a probability of success to clinical trials solely by following drugs through clinical trial phase transitions and comparing intended medical applications. The data for this study was limited to information from a commercial dataset and not available publicly. While an open assessment of all experimental medicines would be preferable, the authors stated “trained analysts would require tens of thousands of hours of labor” (14) to perform such a study using ClinicalTrials.gov, a public source for clinical trials data.

The current lack of public data on successful, failed, and on-going drug studies sparked the development of the Clinical Drug Experience Knowledgebase (CDEK-http://cdek.wustl.edu) with the purpose of creating a public platform to analyze all active pharmaceutical ingredients that have ever been tested in humans, as well as their sponsoring organizations and those participating in pre-approval clinical activites. Based on insights derived from previous studies, we focused on three primary pillars for the first instantiation of CDEK: active pharmacetucial ingredients, organizations, and clinical trials. Each pillar is shown in Figure 1 with surrounding metadata fields. Foreign keys in the database link each pillar together. In the next section, we review the current state of clinical stage pharmaceuticals available in public databases.

**Figure 1:**
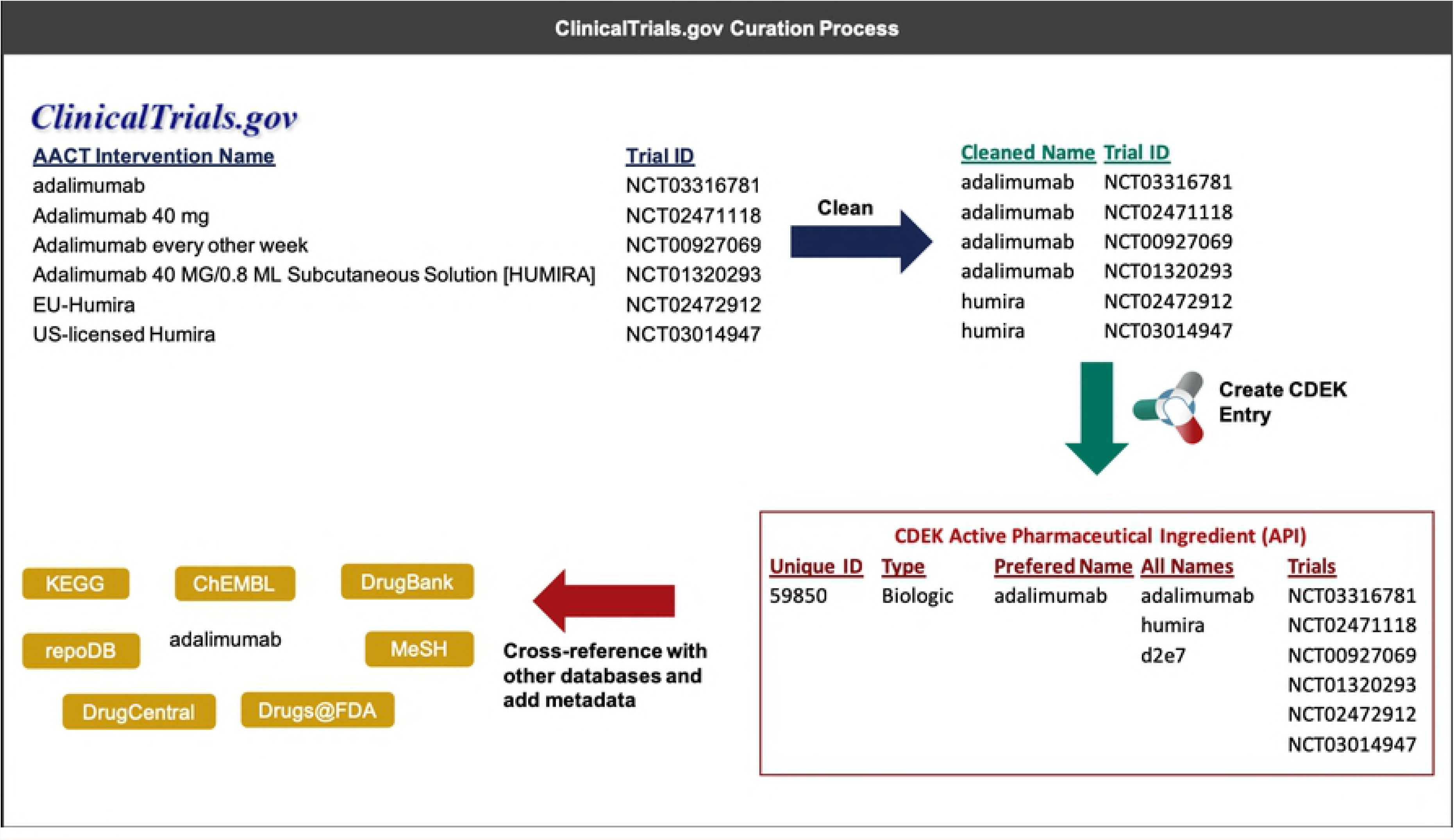
Overview of CDEK contents with three primary pillars: Active Pharmaceutical Ingredients, Organizations, and Clinical Trials. Each metatopic is surrounded with the current fields (solid lines) and planned metadata fields (dashed lines).

### Current state of clinical stage pharmaceuticals in public databases

Several biopharmaceutical databases have emerged over the last decade to enable chemo- and bio-informatics research in the field of drug discovery, including chemical structures to support *in silico* drug discovery, drug repurposing opportunities, and trends in the drug development enterprise. A decade ago, fewer than 200 peer-reviewed articles were published per year referencing a biopharmaceutical database. Today, over 2,500 articles annually cite biopharmaceutical databases and this rate continues to grow exponentially. We recently surveyed several open and freely available databases to explore the current landscape of clinical stage pharmaceuticals and found a collection of databases having drug records that display some evidence of clinical experience.

A selection of databases is listed in Table 1, including a brief description of the clinical content of the database. However, these databases often contain discovery-level or preclinical molecules that have never or will ever enter the clinic. The PubChem (15) database, housing over 100 million compound records, can be filtered to clinical stage compounds by extracting records sourced from ClinicalTrials.gov, ToxCast, or the NCATS Pharmaceutical Collection. ChEMBL (16), another large compound database, can be filtered to clinical stage compounds by selecting records with a max_phase greater or equal to one (with max_phase corresponding to the farthest clinical trial phase the compound has been registered). DrugBank (17), an encyclopedia of active pharmaceutical ingredients, can be filtered to clinical compounds by selecting “Approved”, “Withdrawn”, “Investigational”, “Illicit”, or “Nutraceutical” from their “Drug Group” metadata field. Other databases focus explicitly on approved or withdrawn medicines, making their whole catalog of drugs relevant in terms of clinical experience.

**Table 1:** Public databases containing clinical stage active pharmaceutical ingredients.

| Database | Scope | Clinical Experience Evidence | Access |
| --- | --- | --- | --- |
| **PubChem** | Chemical entities and their bioactivities | Records sourced from Clinicaltrials.gov, ToxCast, or NCATS Pharmaceutical Collection | https://pubchem.ncbi.nlm.nih.gov |
| **ChEMBL** | Bioactivity for drug discovery | Field "max_phase" =>1 | https://www.ebi.ac.uk/chembl |
| **DrugBank** | *in silico* drug discovery and exploration | Field "DRUG GROUP" = "Approved OR Withdrawn OR Investigational OR Illicit OR Nutraceutical" | https://www.drugbank.ca |
| **DrugCentral** | Active pharmaceutical ingredients approved by FDA and other agencies | All records are approved or withdrawn medicines | http://drugcentral.org |
| **SuperDrug2** | Marketed drugs | All records are approved or withdrawn medicines | http://cheminfo.charite.de/superdrug2 |
| **CRIB NME** | FDA approved molecular entities and biopharmaceutical organizations | All records are approved or withdrawn medicines | http://cribdb.wustl.edu |
| **repoDB** | Drug repurposing | All records are either approved or have been in clinical trials. | http://apps.chiragjpgroup.org/repoDB |
| **WITHDRAWN** | Withdrawn or discontinued drugs | All records are withdrawn medicines | http://cheminfo.charite.de/withdrawn |

In a study that inspired the creation of CDEK, our group downloaded the clinical-stage active pharmaceutical ingredients from the sources listed in Table 1. Approximately 11,760 unique active pharmaceutical ingredients with evidence of clinical experience were available collectively from those data sources. However, the total number of active pharmaceutical ingredients that have ever been tested in humans was likely much higher. For example, Wong *et al*. used the Informa Pharma Intelligence databases “TrialTrove” and “Pharmaprojects” to complete their study on estimating clinical trial success rates. In their study, they cited extracting over 21,143 unique compounds from the Informa Pharma Intelligence databases with corresponding clinical trial information (14).

Such findings suggest other active pharmaceutical ingredients may exist in the public domain but have not been curated. ClinicalTrials.gov (accessed through the Aggregate Analysis of Clinical Trials (AACT) database), for example, contains over 286,811 unique trials with over 246,005 unique “intervention names” in a trial (as of 10/20/2018). Multiple “intervention names” correspond to the same active pharmaceutical ingredient. To achieve the ambitious goal of “studying all drugs ever tested in a human”, it was necessary to mine and disambiguate ClinicalTrials.gov data to supplement the compounds available in current open access drug databases.

Descriptions of the disambiguation of ClinicalTrials.gov interventions and organizations follow. Detail on how other databases were used to cross-reference unique ClinicalTrials.gov interventions is also summarized. CDEK is the culmination of this curation effort and is a public database and web platform to interrogate all active pharmaceutical ingredients where there exists objective evidence of human clinical testing. CDEK aggregates metadata surrounding active pharmaceutical ingredients, including the details of clinical trial design, intended indications, and organizations responsible for development. The envisioned use of the CDEK is to support the investigation of many aspects of drug development, including discovery, repurposing opportunities, chemo- and bio-informatics, clinical and translational research, and regulatory sciences. The platform is intended to serve a wide audience interested in investigational agents, which have reached clinical stage development. The uses enabled by CDEK also include the elucidation of broad or focused trends, competitive intelligence, improving drug development efficiency and conveying best practices of lessons learned and future directions.

## METHODS

### CDEK Construction: Curating ClinicalTrials.gov data

Construction of CDEK arose from multiple iterations beginning with the predominant source of data: ClinicalTrials.gov accessed through the Aggregate Analysis of Clinical Trials (AACT) database (18). ClinicalTrials.gov is a repository of clinical trial registrations in the United States and is maintained by the National Library of Medicine (NLM) at the National Institutes of Health (NIH) in collaboration with the Food and Drug Administration (FDA). The AACT database was developed and is maintained by the Clinical Trials Transformation Initiative (CTTI) group, a government-academic collaboration between the FDA and Duke University. The AACT database contains ClinicalTrials.gov data that has been parsed and deposited into a structured relational database. AACT also links clinical trials data to Medical Subject Headings (MeSH terms), a controlled vocabulary containing terms describing disease indications and interventions. This mapping enables querying the data by intervention and disease indication terms. In this first step, we were primarily interested in removing the ambiguity in the trial intervention names and names of sponsoring organizations.

The AACT *interventions* table has the field *intervention_type* with the following distinct terms used to describe an intervention in a trial: Drug, Behavioral, Diagnostic Test, Dietary Supplement, Other, Device, Biological, Procedure, Combination Product, Genetic, and Radiation. To initially populate CDEK with therapeutic clinical trials, all AACT pharmaceutical interventions were included whereas interventions labeled Behavioral, Diagnostic Test, Device, Radiation or Other were excluded. CDEK was populated with associated clinical trial data and organizations linked to those entries. The organizations in turn were parsed from the *sponsors* table*, overall_officials* table, and *responsible_parties* table within AACT. Collectively, these tables contain the lead and collaborating sponsors, trial affiliation data for various study roles (e.g. Principal Investigator, Study Chair), and trial affiliation data for the party type (e.g. Sponsor, Sponsor-Investigator).

In a first round of data cleanup, the names of active pharmaceutical ingredients and organizations were validated. Each active pharmaceutical ingredient was manually labeled by biomedical research curators as being one of either *Vaccine, Gene therapy, Cell therapy, Small molecule, Biologic (synthesized in organisms or cell lines), Biological (derived from human material), Animal product* or *Botanical; and* any active pharmaceutical ingredient not categorized as such was removed from the dataset. Additionally, active pharmaceutical ingredient names were manually curated and any active pharmaceutical ingredient listed as a combination drug was split into its constituent parts. Manual validation and cleaning of active pharmaceutical ingredient names included correcting obvious misspellings and removing salt or solvent forms. Similarly, each organization was labeled as being one of *Individual*, *Academic/Hospital, Government, Foundation, For profit* or *Unknown*, and each organization name was validated and normalized to have consistent naming nomenclature. Figure 2 illustrates an example of the curation process for an active pharmaceutical ingredient.

**Figure 2:**
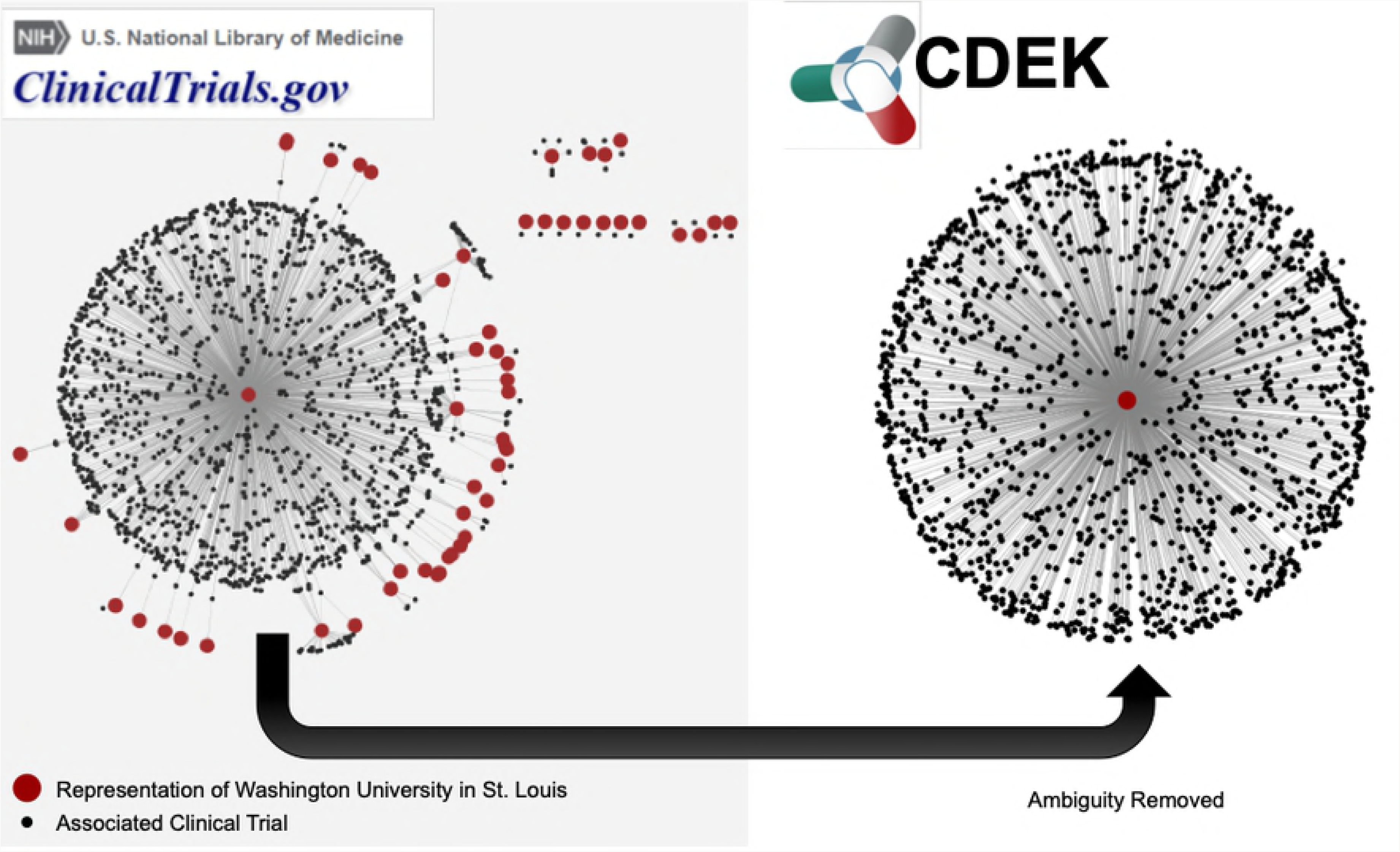
An example that illustrates the process of extracting inteventions from ClinicalTrials.gov (through AACT) and creating a unique active pharmaceutical ingredient record in CDEK. Curation begins by extracting the intervention names from trials containing active pharmaceutical ingredients and cleaning names to strip any perfulous text (e.g. dosing amount, dosing freqency). Once complete, an automated program flags entities that should be merged into a single CDEK record using a set of “merging” criteria. The curation software will also flag entities that are made up of two or more active pharmaceutical ingredients using a set of “splitting” criteria (e.g. the drug “Mavyret” is a combination of two active pharmaceutical ingredients, glecaprevir and pibrentasvir, used to treat hepatitis C). A unique CDEK active pharmaceutical ingredient record is created and assigned a unique id, a type, and a preferred name. All names are stored as synonymns and all trials are linked to the unique active pharmaceutical ingredient ID. Finally, several external databases are cross-referenced to pull metadata and provide hyperlinks to more information about that active pharmaceutical ingredient. This metadata was also used to flag entries that should be merged into a single active pharmaceutical ingredient.

### Construction: Cross-referencing with public biopharmaceutical databases

Additional sources of data were ingested into the database following the first round of cleanup. Several open drug-compound databases containing clinically tested therapeutics to capture active pharmaceutical ingredients with evidence of clinical testing outside of the ClinicalTrials.gov registry. These databases included Drugbank(17), ChEMBL(16), PubChem(15), SuperDrug2(19), DrugCentral(20), WITHDRAWN(21), repoDB (22) and CRIB NME (4). The first three of these databases were subsetted to access only those therapeutics with evidence of clinical testing, while the remainder contain soley clinically-tested therapeutics (approved by a regulatory agency, withdrawn from the market for any reason, or associated with a clinical trial). All DrugBank (v5.0.7) compounds labeled “experimental” were excluded from CDEK as DrugBank defines “experimental” as “drugs that are at the preclinical or animal testing stage.” The ChEMBL database labels drug compound records as having a *max_phase*, the maximum clinical trial phase for which that drug compound has been tested. Any compounds with a *max_phase* greater than 0 was ingested from ChEMBL v23. Any PubChem compound annotated as sourced from ClinicalTrials.gov were ingested. Additionally, all approved drugs listed on the regulatory websites (as of April 2018) of the Food and Drug Administration (Drugs@FDA) and European Medicines Agency (EMA) were parsed, validated and ingested. The metadata provided by these external databases were used to facilitate the disambiguation process described in the next section.

### Construction: Removing ambiguity to get a list of unique Interventions and Organizations

After initial cleanup and ingestion, expert curators split and merged organizations and active pharmaceutical ingredients based on their metadata. We performed this cleanup and ingestion process semi-manually by first programatically flagging data for review followed by manual validation of each flagged entry. The program identified active pharmaceutical ingredients to be considered for merging when two or more distinct entries are were labelled with the same active pharmaceutical ingredient name, *source_api_id* (the ID given to the active pharmaceutical ingredient in a given source), chemical structure (SMILES string), or had overlapping synonyms. Similarly, the program flagged records for splitting active pharmaceutical ingredients into multiple distinct compounds when multiple non-distinct chemical structure data was associated with a given active pharmaceutical ingredient or if multiple *source_api_id*s were associated with the active pharmaceutical ingredient. The program calculated similarity scores (e.g. Levenshtein distance) for all pairs of organizations to identify highly similar organizations pairs, which expert curators then manually validated as either being the same organization or not.

Figure 3 demonstrates an example of the ambiguous nature of ClincalTrials.gov data. Our particular home institution, Washington University in St. Louis (WUSTL), was designated by more than 50 unique representations in ClinicalTrials.gov. This represents the ambiguity challenge to be remedied. Figure 3 shows a network in which all red nodes are different representations of the WUSTL name and all black nodes are the clinical trials associated with that name. After disambiguation, all WUSTL affiliated trials were represented as one organization: “Washington University in St. Louis”. The June 2017 snapshot of AACT has 54047 organization names associated with the 127,220 clinical trials in CDEK. We manually validated and collapsed these entries into 24,728 unique CDEK organizations. Furthermore, AACT has 104,627 unique interventions names that we manually validated and collapsed to 17,096 CDEK active pharmaceutical ingredients. During the curation process, we stored all names, which had been collapsed into single organizations as “alternative names”. This allows for users to search many different terms in our web application.

**Figure 3:**
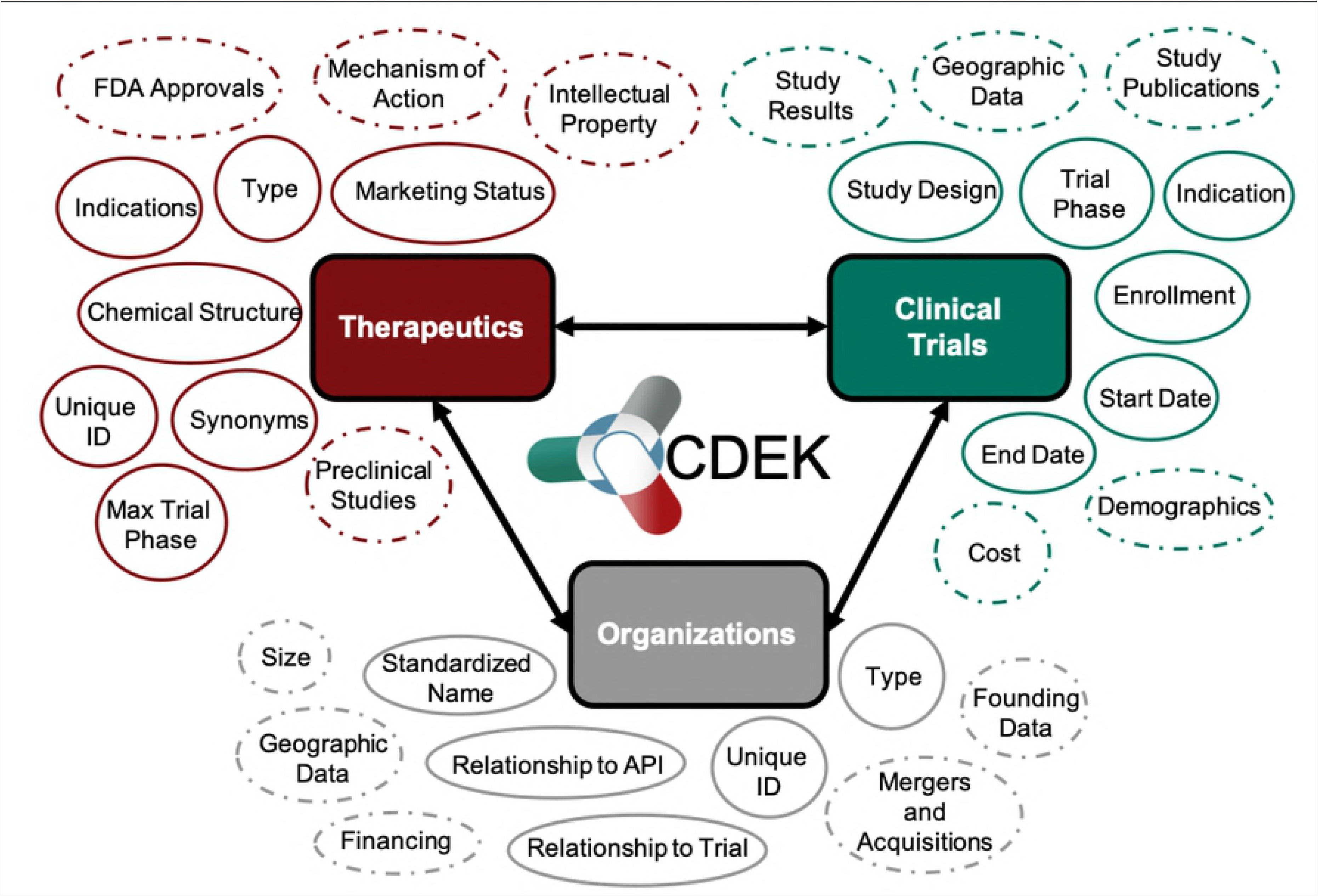
Network graph of trials associated with Washington University in St. Louis. The left graph shows different representations of Washington University in St. Louis in ClinicalTrials.gov as red nodes. Examples of different names representing “Washington University in St. Louis” include: “Washington University School of Medicine”, “Washington Universite Siteman Cancer Center”, and various misspellings of the word ‘university’. Black nodes are the clinical trials associated with each different name for the Washington University in St. Louis organization. The right graph shows CDEK data with Washington University in St. Louis as a single organization with its corresponding clinical trials.

### CDEK Contents

Table 2 provides summary statistics of CDEK contents: active pharmaceutical ingredients (n = 22,292), clinical trials (n = 127,223), and organizations (n = 24,728).

**Table 2:** Summary counts of CDEK data

| Organization Type | Count | API Type | Count | Trial Phase | Count |
| --- | --- | --- | --- | --- | --- |
| Academic/Hospital | 9495 | Small Molecules | 13169 | Phase 2 | 32538 |
| For Profit | 6577 | Biologics | 2583 | Phase 1 | 23656 |
| Individual | 3634 | Botanicals | 1769 | Phase 3 | 22641 |
| Unknown | 3183 | Vaccine | 1698 | N/A | 18830 |
| Foundation | 1200 | Cell Therapy | 1521 | Phase 4 | 18267 |
| Government | 658 | Biological | 1182 | Phase 1/Phase 2 | 7054 |
| **Total Orgs** | **24747** | Animal Product | 233 | Phases 2/Phase 3 | 3184 |
|  |  | Gene Therapy | 157 | Early Phase 1 | 1163 |
|  |  | **Total APIs** | **22312** | **Total Trials** | **127333** |

CDEK includes all prophylactic and therapeutic chemical or biological entities, including but not limited to vaccines, cell therapies, gene therapies, animal products, and biologics – many of which are not typically included in other popular compound-oriented databases.

## RESULTS

### CDEK Platform

The CDEK platform used the open-source web framework, Django, which follows the model-view-controller architectural pattern. This allows the internal representation of data (the models) to be separated from the presentation to the end user (the view). In the back-end, the models were implemented as a PostgreSQL database and all data is hosted on Heroku. The controller and views rendered the front-end of the platform using a mix of HTML, CSS and javascript.

The CDEK platform provided two query functionalities, allowing users to quickly interface with the data without having any prior familiarity with a structured query language (SQL). The first functionality, a *basic search* (http://cdek.wustl.edu/search/) enable the user to do a fuzzy, case-insensitive search for keywords or synonyms in order to find either active pharmaceutical ingredients or organizations. This functionality serves as a quick, simplified means of interacting with a single datum. The result displays summary statistics of the basic CDEK pillars. Figure 4 shows an example of active pharmaceutical ingredients and organization summary pages. For an active pharmaceutical ingredient, the clinical trial distribution is plotted according to trial phase and organizations involved in its developed is plotted according to organization type. For an organization, the involvement in clinical trials and active pharmaceutical ingredient development is plotted according to trial phase and active pharmaceutical ingredient type, respectively. In both search displays, a list of alternative names is given. For those interested in the source data, or who seek to visualize the ingested reference, CDEK allows the user to link to external cross-referencing databases.

**Figure 4:**
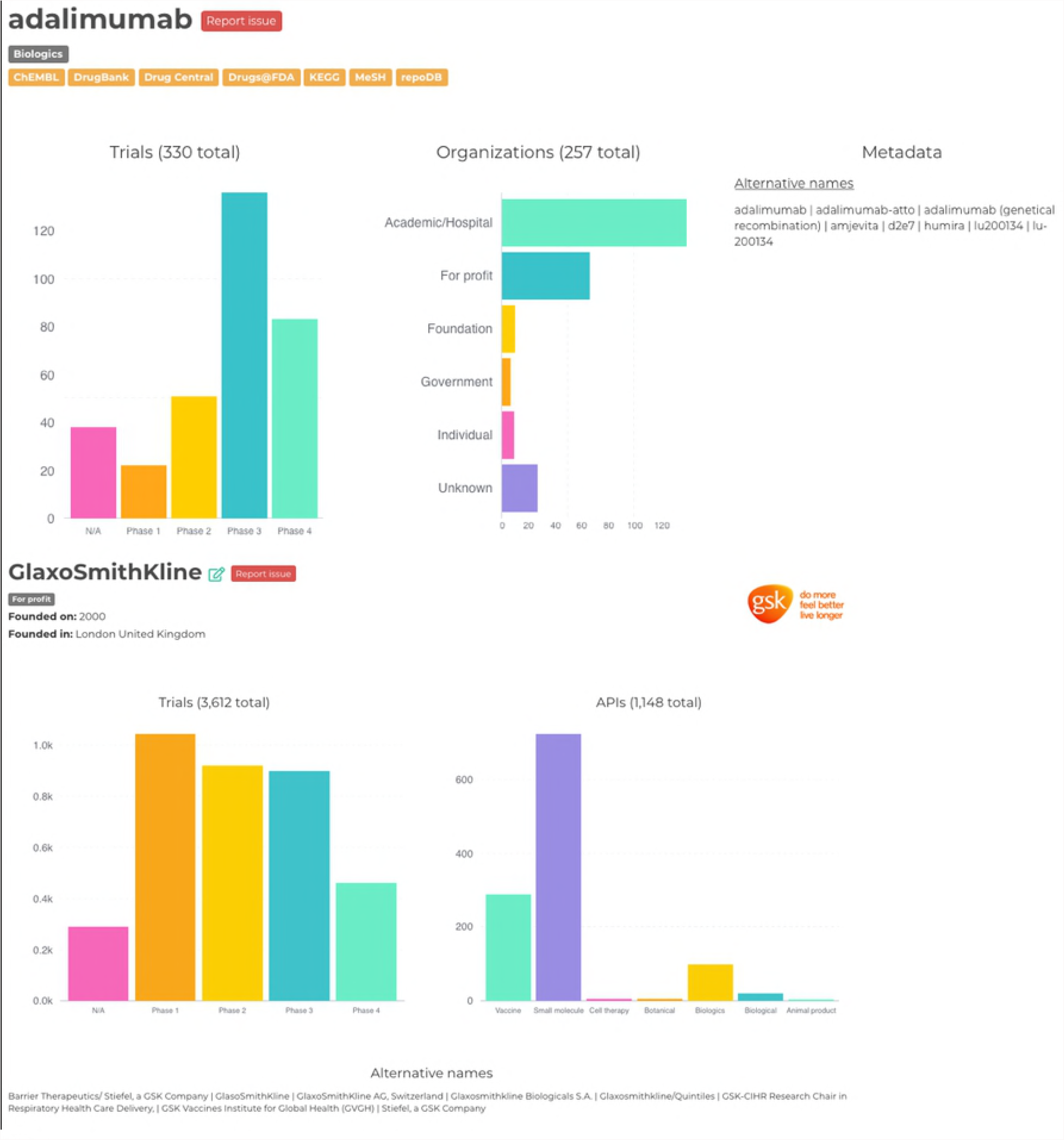
Example Active Pharmaceutical Ingredient and Organization summary pages from the CDEK platform. Adalimumab, was the top selling drug of 2017 while GlaxoSmithKline has the most associated clinical trials in the CDEK database.

Users are directed to an advanced query functionality to access the granular CDEK data.

The *advanced query* functionality (cdek.wustl.edu/query/) provides users with more control over the metadata are used to filter the dataset. A dynamically generated user-interface (UI) allows a user to build a SQL-like query, in a *WISYWIG* (“what you see is what you get”) fashion, without having any previous knowledge of SQL. Complex queries can be quickly generated by building filtration rules (predicates) and by combining them with boolean logic. These data are then submitted to the back-end through an AJAX (Asynchronous JavaScript And XML) call to a database-view which combines all the CDEK data into a single table. This AJAX call initializes a Celery worker which will process the query request on a separate Heroku worker dyno and return the result in a non-blocking fashion; this ensures that the platform can scale properly as more queries are submitted and ensures a better user experience. Results are presented in a familiar table-like manner with sortable columns and hyperlinks to individual data instances. A RESTful API (application programming interface) provides an endpoint for viewing these individual data when either requesting a single active pharmaceutical ingredient or organization instance. This endpoint dynamically generates interactive charts which summarize the data for the given data instance. Our advanced query builder allows a user to filter CDEK data to granular details. Figure 5 shows a screen shot of the query tool web application. In this example, the data returned will be all unique Phase III clinical trials (n = 681) studying lung or cardiovascular diseases, excluding vaccines, and run by GlaxoSmithKline as the lead sponsor between 2012 and 2017.

**Figure 5:**
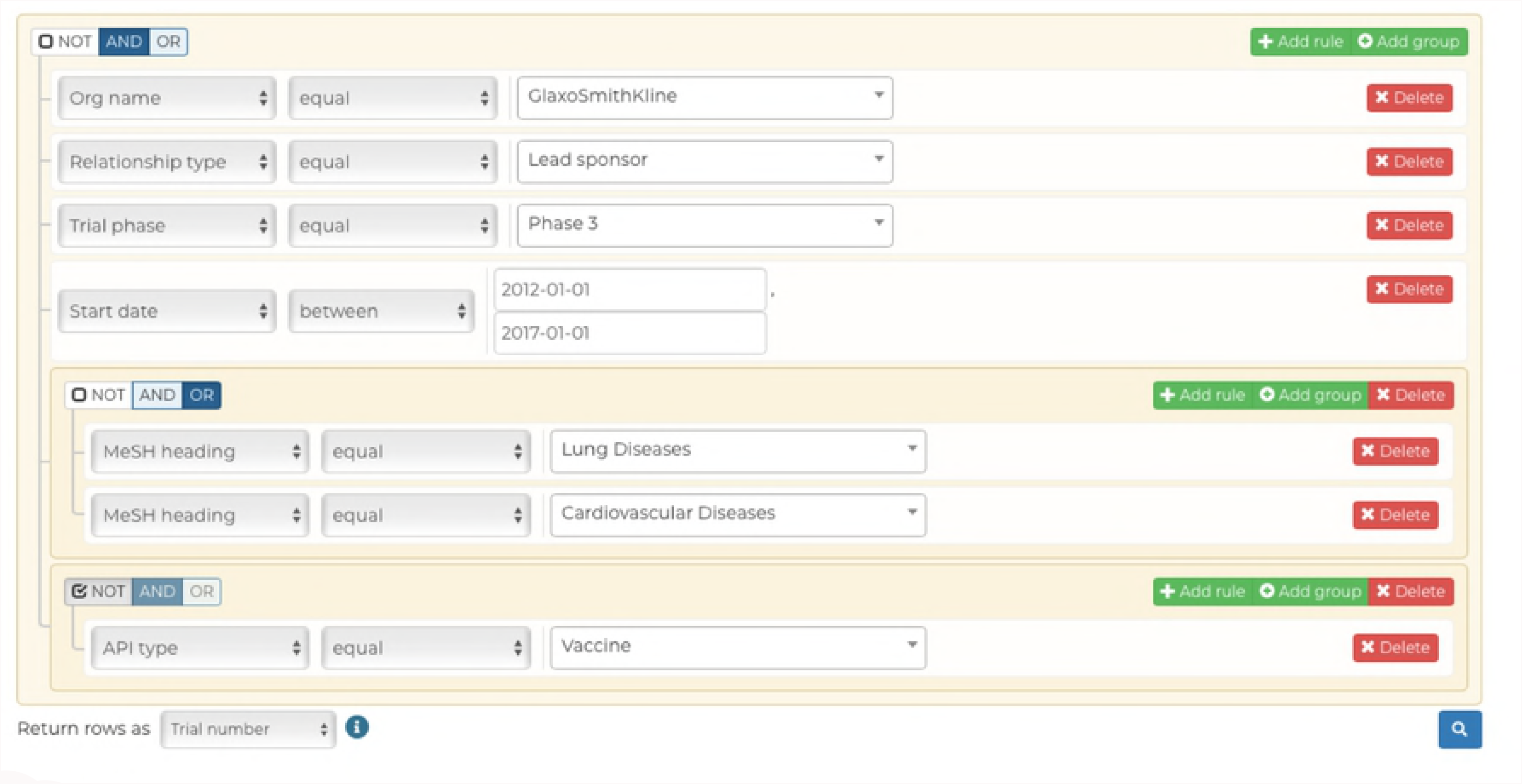
Our advanced query builder allows users to filter down CDEK data to very granular details. In this example, the data returned will be all unique Phase III clinical trials studying lung or cardiovascular diseases, excluding vaccines, that were ran by GlaxoSmithKline as the lead sponsor between 2012 and 2017.

### Lessons Learned

Approximately 17,096 unique active pharmaceutical ingredients in CDEK were sourced from ClinicalTrials.gov, 9,781 of which currently cannot be found in any databases cross-referenced in CDEK (see Table 1). These active pharmaceutical ingredients comprise 3160 small molecules, 1477 vaccines, 1438 cell therapies, 1387 biologics, 1084 botanicals, 982 biologicals, 143 gene therapies, and 110 animal products. The databases included for initial cross-referencing primarily focus on small molecules and biologics. Therefore, we reviewed unique small molecules and biologics extracted from ClinicalTrials.gov, hereafer refered to as “unique CDEK records”. Most (90%) unique CDEK records have been registered in three or fewer clinical trials and 85% of the clinical trials referencing these drugs are prior to Phase III. This indicates that early stage active pharmaceutical ingredients might not typically be flagged for curation in traditional databases. Another interesting trend is almost two-thirds (64%) of the unique CDEK records were sponsored by for profit organizations. This contrasts to the whole CDEK dataset where less than one third (30%) of all trial lead sponsors are for profit organizations.

The active pharmaceutical ingredient contents of CDEK was compared with other common compound-oriented drug databases including: PubChem, Chembl, DrugBank, DrugCentral, SuperDrug2, WITHDRAWN, repoDB, and drugs@FDA. Despite our initial assumption that existing databases, once aggregated, would convey a comprehensive list of experimental medicines, approximately 43% of active pharmaceutical ingredients in the CDEK database were extracted from AACT and cannot be found in any of the other compound-oriented databases listed above.

We reviewed the overlap of active pharmaceutical ingredients with evidence of clinical testing among several open databases, including those listed in Table 1, AACT, and the Drugs@FDA database. Figure 6 shows the this overlap as a heatmap, comparing content across several drug databases. This visualization demonstrates that some databases are almost complete subsets of others (99% of repoDB compounds can be found in ChEMBL, DrugCentral and DrugBank). PubChem, one of the largest compound libraries showed consistently high overlap values across the spectrum. The overlap between AACT active pharmaceutical ingredients and PubChem is the highest, closely followed by AACT and ChEMBL.

**Figure 6:**
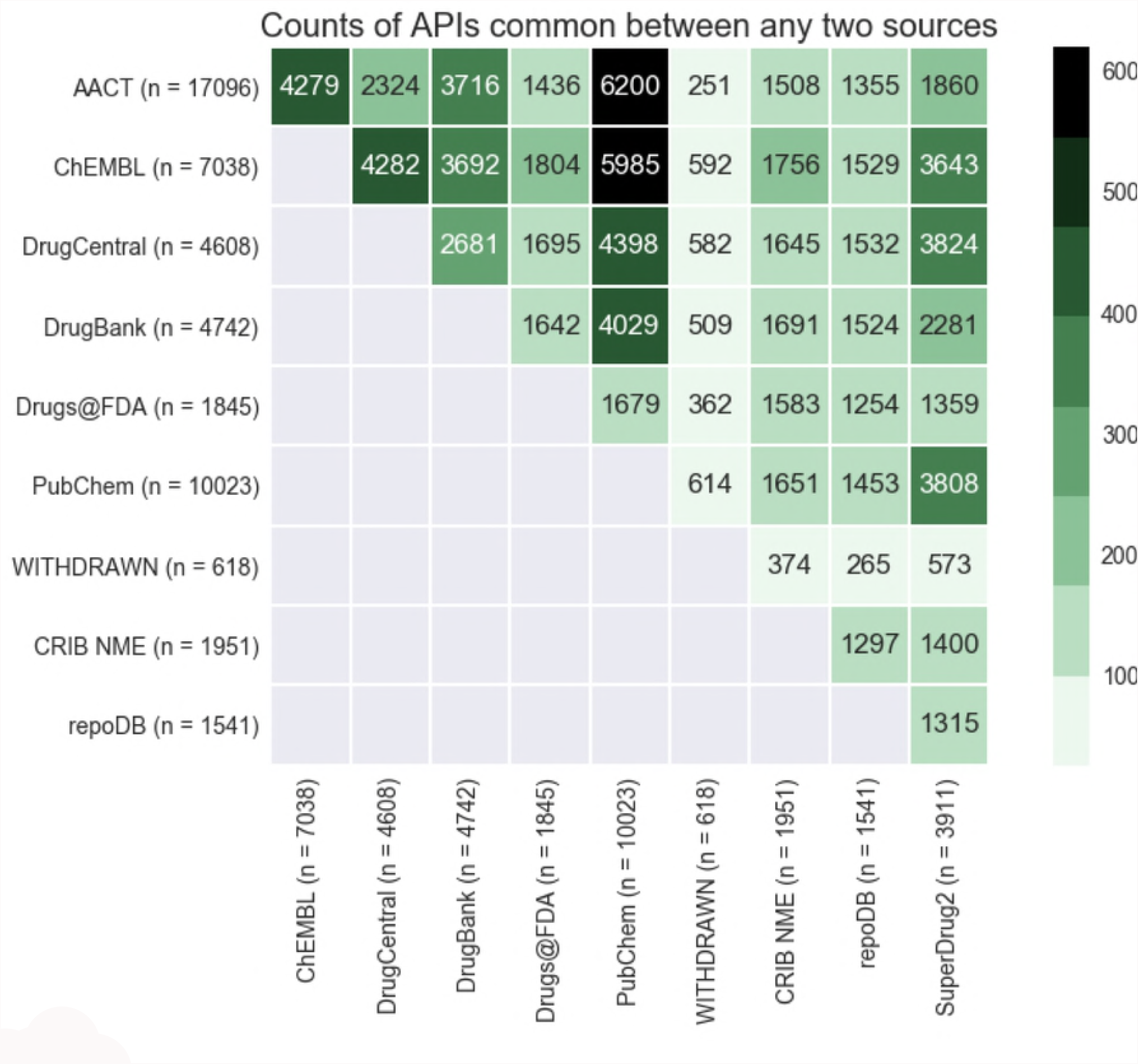
Heatmap displaying the overlap in active pharmaceutical ingredients (APIs) between any two databases in CDEK. The coloring and number displayed at the intersection between any two databases is the total number of shared APIs. The total number of unique APIs from each database that has evidence of clinical experience is noted in paranthesis next to each database name label.

## DISCUSSION

The purpose of CDEK is to provide researchers with an open database and platform to study the entire drug development enterprise by interrogating *all* active pharmaceuticals with evidence of clinical testing. While not truly comprehensive, we have created the first release of such a resource and below we discuss several on-going strategies for improvement.

The first instantiation of CDEK was derived from a June 2017 snapshot of the AACT database. Over 20,000 trials registered in ClinicalTrials.gov were not included in the first instantiation but we are currently developing a novel “ingestion pipeline” to allow curators to update the data automatically and in real time. Databases listed as cross-referencing sources will be updated in CDEK in the future along with the addition of new data sources – such as ToxCast and ZINC. Future curated databases will also be merged into CDEK under the conditions they are public, verifiable and contain evidence of clinical-trial candidates.

The curation of several new metadata fields will be incorporated into CDEK. These fields are summarized in Figure 1 encircled by dashed lines. They include information such as patents surrounding active pharmaceutical ingredients, approval status of each indication associated with an active pharmaceutical ingredient, clinical trial study results, and the merger and acquisition activity of for-profit organizations conducting clinical trials.

Another on-going area of development is mining scientific publications containing clinical trial information. ClinicalTrials.gov was created in response to the Food and Drug Administration Modernization Act of 1997 (FDAMA), with the first public version of ClinicalTrials.gov released in 2000. Therefore, it is necessary to search public reports of clinical studies for trials that may not have been registered, or that were conducted prior to 1997.

Finally, continued efforts are being made to clean and disambiguate any residual errors propagated through the initial data cleanup. We intend to employ higher standards for chemical data set curation methods, such as those outlined by Fourches et al (23). Due to the expansive efforts needed to keep CDEK up-to-date and accurate, our group is also interested in deploying crowd-based curation methods in the future.

## CONTACTING CDEK

CDEK was developed and is maintained by the Center for Research Innovation in Biotechnology (CRIB) at Washington University in St. Louis. CRIB studies the blend of science, business, and regulation of biotechnology, medical devices, and healthcare IT to ensure continued improvements in the delivery of medical innovations and public health. CRIB is actively pursuing collaborations to study the data within CDEK. Errors and suggestions for improvement can be submitted at http://cdek.wustl.edu/about/. Or contact us via e-mail at cdek at wustl dot edu.

## ACKNOWLEDGEMENTS

The authors would like to acknowledge Tom Krenning for the initial conceptualizations of CDEK. Research reported in this publication was supported by the Washington University Institute of Clinical and Translational Sciences grant UL1TR002345, sub-award TL1TR002344, from the National Center for Advancing Translational Sciences (NCATS) of the National Institutes of Health (NIH). The content is solely the responsibility of the authors and does not necessarily represent the official view of the NIH.

